# Discovery of a locus associated with susceptibility to esca dieback in grapevine

**DOI:** 10.1101/2023.10.20.563213

**Authors:** Arnold Guillaume, Prado Emilce, Dumas Vincent, Butterlin Gisèle, Duchêne Eric, Avia Komlan, Merdinoglu Didier

**Affiliations:** INRAE, Université de Strasbourg, UMR SVQV, Colmar, France

## Abstract

Esca is the most destructive and predominant grapevine trunk diseases. The chronic infections and vine mortality caused by esca syndrome lead to huge economic losses and threatens the sustainability of vineyards worldwide. Although shown as associated with the presence of wood fungi, the etiology of esca remains still unclear and putatively involves multifactorial causes, which makes the development of effective control methods challenging. As differences in esca susceptibility had already been observed among grapevine varieties, we investigated in a biparental population the presence of genetic factors that can explain theses variations. Thanks to the destructive phenotyping of a 16-year-old vineyard plot, we discovered that the Gewurztraminer variety carries on chromosome 1 a locus linked to variations in trunk necrosis associated with esca, which we have named *ENS1*. Our study also suggests that there is a partial link between trunk vigor and necrosis due to esca. To our best knowledge, *ENS1* is the first instance of genetic factor identified as involved in the limitation of necrosis associated to grapevine esca. While the identification of *ENS1* alone may not provide a complete resolution of the esca issue, this discovery represents nonetheless a first step towards a genetic solution and paves the way for broader genetic investigations in the future.

## Introduction

Grapevine (*Vitis vinifera* L.) is among the most important perennial crops, not least thanks to its economic weight and its role in shaping the landscape^1–4^. Nevertheless, vineyards are affected worldwide by many severe diseases, which negatively impact berry quality, plant growth and yield, even leading to the death of infected plants. Among them, grapevine trunk diseases (GTDs), which includes eutypa dieback, esca, and botryosphaeria dieback, threaten the sustainability of viticulture worldwide and are considered the most destructive diseases of grapevine for the past three decades ^5,6^. GTDs are associated to the presence of fungi which colonize the permanent woody structure of grapevines, causing chronic infections^7^. Globally, the economic cost of grapevine replacement, required because of the mortality, is over $1.5 billion per year^8^. Since the last few decades, the incidence of GTDs has rapidly increased in all wine-producing countries. In Spain, for example, the percentage of affected vines increased from 1.8% in 2003 to 10.5% in 2007^6^. In France, a six-year survey showed that the esca and botryosphaeria dieback incidence increased sharply between 2003 and 2008 in several major wine-producing regions, reaching 11% for the most affected of them^9^.

In established vines, esca is the most destructive and predominant GTD^8,10–12^. The fungi associated with the esca syndrome are primarily the ascomycetes *Phaeoacremonium spp*. and *Phaeomoniella chlamydospora*, and the basidiomycete *Fomitiporia mediterranea*^13^. It is hypothesized that they act in sequence, *Phaeoacremonium aleophilum*. and *P. chlamydospora* colonizing wood first^14^. However, the etiology of esca is still unclear and several multifactorial scenarios are under consideration to explain the expression of symptoms^14–17^. Indeed, in addition to biotic agents, some scenarios involve abiotic factors, in particular, those leading to high vigor^18–20^. The influence of climate change in favor of disease expression has also been suggested recently^21^.

Esca is characterized by so-called “tigerstriped” leaf symptoms and by the development of various internal necrosis in wood tissues^17^, mainly degraded wood and white rot tissue^10,12,22^. Wood necrosis can be the result of wounds that cause healing cones but also of the degradation of living wood by *P. chlamydospora* and *P. aleophilum*^23^. White rot is an evolution of degraded wood mainly caused by *F. mediterranea*^24^, the most common saprophyte associated with affected tissues and considered as the main agent within the esca disease complex^25^. Observations on cross sections of trunks showed that the greater the extent of necrosis, the higher the mortality rate of the plants and that white rot extent is positively correlated with the total necrotic area of a trunk^26^. The link between the extent of internal necrosis and the severity of foliar disease symptoms was documented by several reports^27,28^. Two recent studies have provided new evidence of the relationship between leaf symptoms and wood necrosis. Ouadi *et al*.^29^ observed that, in plants with leaf symptoms, at least, 10% of wood had been affected by white rot. Moreover, by integrating the history of foliar symptom expression over years, Fernandez *et al*.^22^ reported a strong correlation between wood necroses and foliar symptoms.

Since the ban of sodium arsenate, which was the only curative chemical against esca, the currently proposed methods to fight against GTDs are mostly preventive and aim at mitigating the disease effect. Producing healthy plants in nurseries thanks to wound protection and hot water treatments, applying prophylactic measures that limit the spread of inoculum in the vineyards, practicing a training system which avoids big wound or allows trunk renewal are considered effective to slow down the disease propagation^6^. However, the main practical measures currently used to control esca in the vineyard aim at limiting the necroses, either at prophylactic level, through pruning to limit the formation of dead wood, or thanks to trunk surgery, which consists in removing white rot inside the trunk^30^, and allow to significantly cure symptomatic vines^31^.

Besides preventive and curative methods, genetic diversity of susceptibility to GTDs has also been explored to identify species, varieties or clones which could be used through a breeding strategy^9,32–34^. Observations of esca susceptibility of grapevine varieties evaluated in the vineyard in independent experimental settings are often consistent with each other. For example, in three studies conducted in Italy, the esca incidence recorded on Chardonnay was invariably low, whereas Sangiovese and Trebbiano were moderately affected and Cabernet Sauvignon was the most severely attacked^32–34^. It is thus safe to assert that the expression of esca symptoms in the vineyard is partly linked to the genetic nature of the plant material^9^. However, discrepancies are also observed between studies, suggesting that other factors such as rootstock, soil or weather conditions modulate the disease expression^9,34^.

Although genetic resistance bears promise as a tool to reduce the incidence of esca and other GTDs, no genetic study has, to date, allowed to identify a genetic factor that can explain the variations in susceptibility observed between grape cultivars in the vineyards. In our present work, we addressed this issue thanks to progenies derived from two varieties, Riesling and Gewurztraminer, considered as different for their susceptibility to esca. Because our aim was to observe wood symptoms recognized as related to the severity of the disease under production vineyard conditions, we have assessed the extent of internal trunk necroses on mature vines thanks to destructive longitudinal sections. Resulting genetic analyses allowed us to identify and locate a locus linked to variations in trunk necrosis associated with esca.

## Materials and methods

### Plant material

Our study focused on Riesling and Gewurztraminer grape varieties and their progeny. These two grape varieties are susceptible to esca, although the susceptibility is generally more pronounced for Gewurztraminer, according to the observations made in the Alsatian vineyards^35^.

The observations were performed on two different populations planted in two distinct experimental designs, as follow:

□ Experiment A: Riesling x Gewurztraminer (RIXGW). Located in the Alsatian PDO vineyard (Bergheim, France), the experiment included 382 genotypes, progeny of a cross between Riesling clone 49 (RI) and Gewurztraminer clone 643 (GW), with elementary plots consisting of three plants per genotype. The control modalities were represented by 12 elementary plots of Riesling clone 49 (RI) and 13 of Gewurztraminer clone 643 (GW) evenly distributed throughout the experiment. The experiment was planted in 2006 with plants grafted on rootstock 161-49C clone 198.
□ Experiment B: S1 Gewurztraminer(S1GW). Located in Colmar (France), outside the Alsace PDO, the experiment included 90 descendants of Gewurztraminer self-fertilization, with elementary plots consisting of three plants per descendant. The control modality was represented by 11 elementary plots of GW distributed throughout the experiment. The experiment was planted in 2002 with plants grafted on grafted on rootstock 161-49C clone 198.

For both experiments, vines were trained with a double Guyot system on a vertical trellis at a planting density of 4800 plants per ha.

### Phenotyping

#### Longitudinal sections of the trunk

In March 2022, each stump was cut vertically down with a chainsaw, starting from the head of the plant to 5 cm below the grafting point. The half-part detached from the rootstock was discarded. The other half, remained attached to the rootstock, was photographed with a uniform blue background and a 100 cm^2^ reference area (Figure S2).

#### Visual scoring of necrosis

After cutting, a visual estimation of the presence of white rot was made in situ using a notation scale (V_WR) ranging from 0, meaning absence of white rot tissue, to 10, meaning more than 80 % of white rot tissue (Figure 1A). A visual estimation of the presence of the total necrosis, *ie*. sum of degraded wood and white rot tissue, (V_TN) was also performed with the same notation scale (Figure 1B).

**Figure 1.**
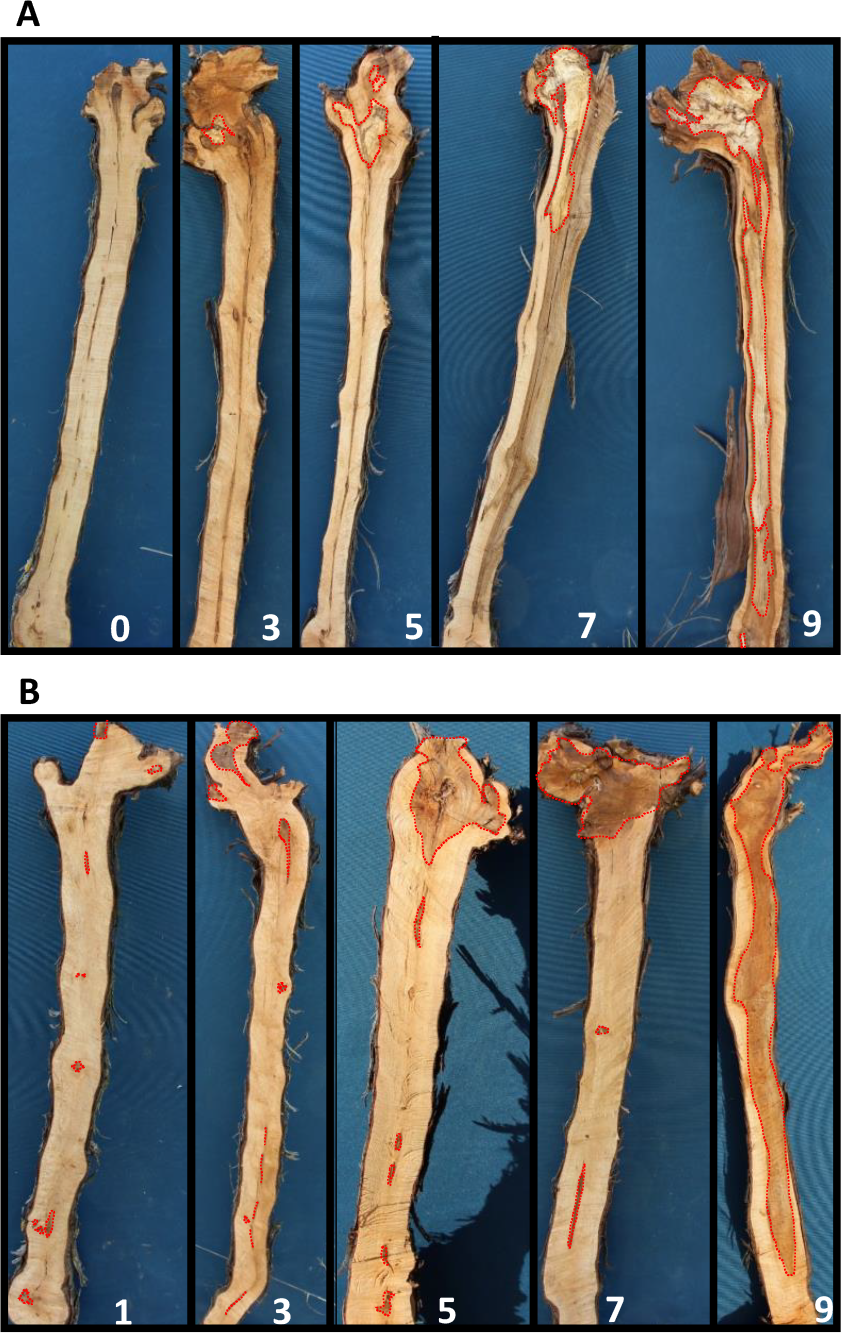
Notation scales for white rot and total necrosis. A visual estimation of the importance of necrosis was made in situ on trunk sections using to a notation scale ranging from 0, meaning absence of necrosis, to 10, meaning more than 80 % of necrosis for both white rot (V_WR; examples on panel A) and total necrosis (V_TN; examples on panel B). The observed necroses are surrounded by a red dotted line.

#### Image analysis

Image analysis was only performed on experiment A.

The following steps were applied to each picture (Figure S3):

▫ homogenization of the background by replacing it by a uniform green were treated with Gimp v2.10.30;
▫ delimitation by hand of white rot tissue and uniformly coloring it;
▫ separation of the various image components - background, reference surface, white rot, healthy wood, degraded wood and bark - with the pixel classification function of Ilastik v1.3.3, after training on a set of training pictures;
▫ recovering the number of pixels of each of the 6 components with Image J 1.53r;
▫ calculation of each variable according to the following formulas, with C1 class corresponding to background, C2 to reference surface, C3 to white rot, C4 to healthy wood, C5 to degraded wood and C6 to bark: I_TA = (C3+C4+C5) * 100/C2, for the area of the trunk section in cm^2^); I_WR = C3*100/(C3+C4+C5), for the proportion of white rot in % of trunk section area; I_TN = (C3+C5)*100/(C3+C4+C5), for the proportion of total necrosis in % of trunk section area.

### Statistical analyses

All the statistical analyses were performed with the R Statistical Software (v4.2.2). Briefly, we used the `kruskal.test` function for Kruskal Wallis tests, `cor.test` function Pearson correlation, the `car` package for calculation of heritabilities and covariance analysis and `ggplot2` package for plotting graphs. Broad-sense heritability was calculated as H^2^ = (σ^t2^-σ^e2^)/σ^t2^, where σ^t2^ is the total variance observed over all the genotypes from the RIxGW progeny and σ^e2^ the residual variance extracted from the anova of the RI and GW elementary plots in experiment A.

### Genetic analysis

#### DNA extraction

Total DNA was extracted from 80 mg of young expanding leaves using a DNeasy^®^ Plant Mini Kit (Qiagen S.A., Courtaboeuf, France) as described by the supplier.

#### Genotyping-by-sequencing approach and construction of genetic maps

QTL mapping was carried out using 252 genotypes from the RIxGW progeny defined as reference population. Genetic markers consisting of single nucleotide polymorphisms (SNPs) were obtained by a genotyping by sequencing (GBS) according to Elshire *et al*.^40^, and modified as described in Chedid^41^.

The final library was sequenced on Illumina HiSeq 2000 platform (paired-end 2 x 100 bp). Raw reads were cleaned and trimmed with cutadapt (version 3.7)^42^ and then aligned on the grapevine reference genome PN.v4^43^ using BWA (version 0.7.17)^44^. Individual bam files were filtered with samtools (version 1.15.1)^45^ and then fetched into Stacks v2.60^46^, using the modules “gstacks” and “populations” to produce a vcf file containing the variants detected across individuals. The obtained vcf file was then filtered using bcftools (version 1.9)^45^, removing loci with average missing data > 10%. Samples with the remaining loci had a maximum of 30% of missing data (and only 18 samples had a missing data level > 10%, among which only 5 had a level > 20%) and were all kept for the subsequent steps.

The two parental genetic maps were built using Lepmap3^47^. Briefly, the final vcf file was submitted to the module ParentCall2 together with a pedigree file, to call segregating markers. The call file went then through the module SeperateChromosomes2 to split the markers over linkage groups. Nineteen linkage groups with confident support were retained and the markers were ordered on each linkage group using the module OrderMarkers2. Thirty runs were performed for each linkage group and the best run based on likelihoods was retained for each. For parental maps, the option “informativeMask=13” or “informativeMask=23” in the module SeperateChromosomes2. The parameter « grandparentPhase=1 » in module OrderMarkers2 allowed to obtain phased data that was converted to fully informative “genotype” data by the script map2gentypes.awk. The parental genotypes are always “1 2” and the data is phased so that the first digit of the genotypes of the progeny is inherited from the male parent and the second from the female parent. This means that the raw genotypes of the progeny after the conversion were “1 1”, “1 2”, “2 1”, or “2 2” with the first digit inherited from the GW (male) parent. The parental genetic maps are equivalent to a backcross type map. Consequently, for the GW map, we consider the genotype of the female RI parent as always non informative homozygote “1 1” and the progeny genotype categories “1 2” and “2 2” modified to “1 1” and “2 1” respectively. Likewise, for the RI female map, the progeny genotype categories “2 1” and “2 2” were modified to “1 1” and “1 2” respectively. For both maps, the expected progeny genotypes were therefore either homozygote “1 1” (equivalent to AA genotype and called A for simplicity) or heterozygote “1 2” or “2 1” (equivalent to AB genotype and called H for simplicity). Under such a backcross like setup, we may detect QTL only if the A allele is not dominant. Genotypes A are therefore considered homozygous recessive.

#### QTL detection

QTL detection was performed using the R package R/qtl^48^. Briefly, one-dimension scanning was performed using the `scanone` function with the Haley-Knott regression. QTL significance thresholds at p=0.05 were obtained with 1000 permutations. The percentage of variance explained by a QTL was assessed with analysis of variance using type III sums of squares using the `fitqtl` function. Confidence intervals were calculated as Bayesian credible intervals using `bayesesint` function with a probability of coverage of 0.95.

#### Design and genotyping of the Chr1_10021151 KASP marker

Sequences of Gewurztraminer clone 643, available in our laboratory, were analyzed to design a set of primers suitable for KASP analysis in the interval chr1:10021151-10021450 flanking a SNP (position chr01:10029666 of 12X.v2 reference genome assembly) located in the *ENS1* region (Table S4).

Genotyping of the SNP was performed in simplex by the Gentyane platform ((INRAE, Clermont-Ferrand, France) using KASPar chemistry (LGC Genomics, KBS-1016-017; https://gentyane.clermont.inrae.fr/uploads/files/Gentyane_services_v1.pdf).

## Results

### Development of esca-associated necroses differs between Riesling and Gewurztraminer and segregates in their progeny

Riesling (RI), Gewurztraminer (GW) and their progeny from a RI x GW cross (RIxGW; 382 descendants) were planted in an experimental plot in the vineyard (experiment A: elementary plot of 3 plants (e.p.); 1 e.p. for each RIxGW descendant; 12 e.p. of RI and 13 e.p. of GW). This population showed signs of decline 16 years after plantation and we decided to use it to analyze its susceptibility to esca necrosis.

Total necrosis (TN) and white rot (WR) development were recorded after longitudinal section of each plant trunk. The proportion of the section area affected by necrosis was then assessed thanks to two methods: i) by visual estimation (scoring done on a scale of 0, meaning absence, to 10, meaning more than 80 % of affected tissue; V_WR for white rot tissue; V_TN for total necrosis, ie. sum of degraded wood and white rot tissue; Figure 1); ii) by image analysis (I_WR for proportion of white rot in % of trunk section area; I_TN for proportion of total necrosis in % of trunk section area). Image analysis was also used to measure the longitudinal cross-sectional area of the trunk (I_TA for in cm^2^).

RI and GW parents were significantly different for the proportion of trunk presenting necrosis (V_TN) (Table S1). Gewurztraminer was the most affected by esca, with total necrosis. This trend was also observed for the other necrosis variables (V_WR, I_WR and I_TN), although statistically not significant. Trunk development (I_TA) of Gewurztraminer was significantly higher than of Riesling.

RIxGW progeny segregated for all the measured traits and displayed transgressive phenotypes compared to its parents, RI and GW (Figure 2). Total necrosis ranged from 1 to 8.3 for visual scoring and from 4.7 to 41.6 % for the imaging method. White rot was between 0 and 5.7 for visual scoring and between 0 and 16.2% for imaging method. The surface of the trunk section assessed by image analysis also varied greatly, from 201.3 to 505.9 cm^2^ (Table 1). In order to estimate the part of the variation due to genetic effects, braod-sense heritability was calculated for each variable. Overall, the values were moderate to high, ranging from 0.240 for V_TN to 0.563 for I_TA. Visual scoring gave lower heritabilities than image analysis for total necrosis and white rot (Table 1).

**Figure 2.**
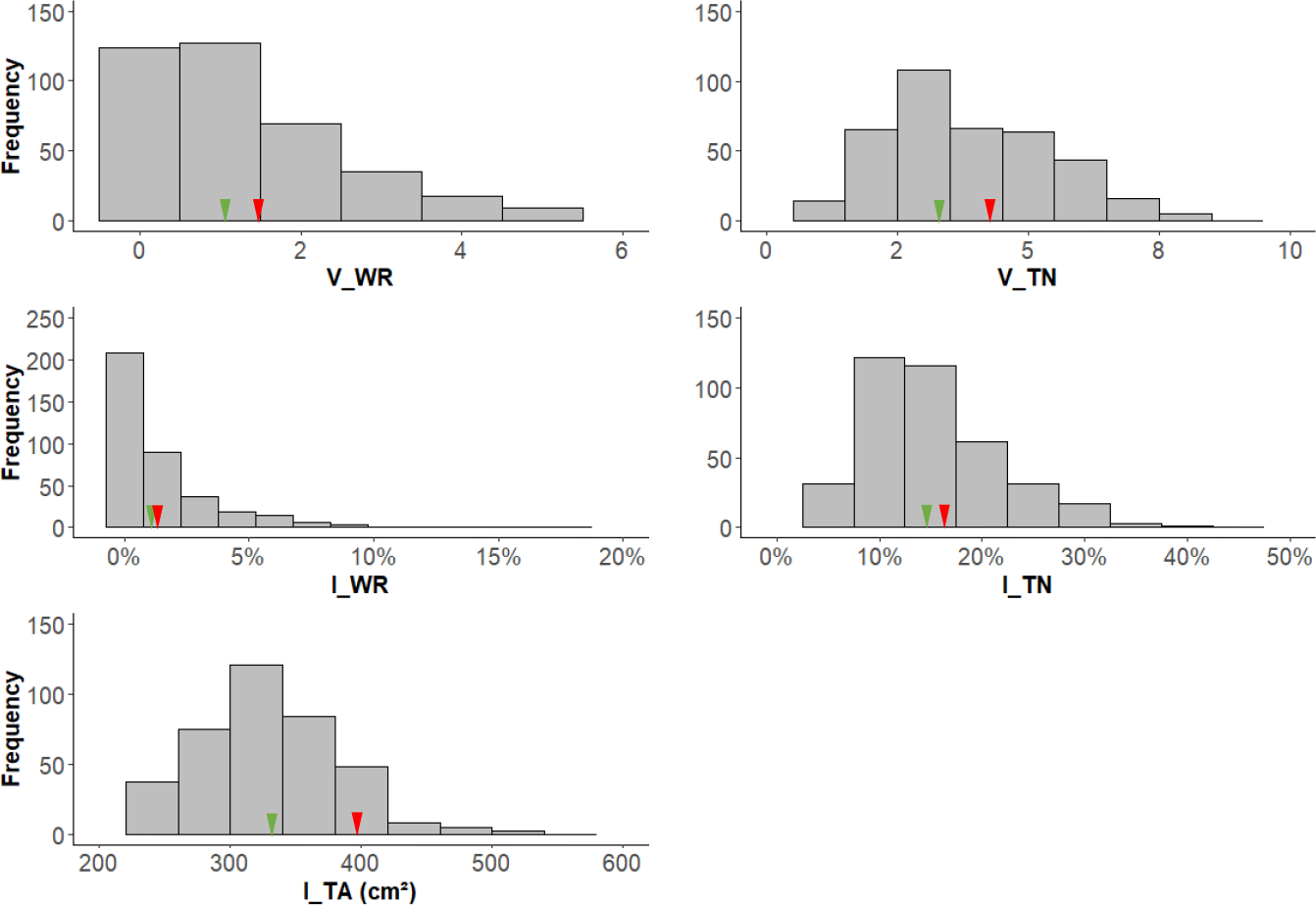
Distribution of necrosis and trunk vigor parameters in the RIxGW progeny. Mean values for RI and GW controls are represented with green and red arrows, respectively.

**Table 1.** Descriptive statistics and broad-sense heritabilities of RIxGW population. The calculations were made from the data recorded in experiment A. σ^t2^ is the total variance observed over all the genotypes from the RIxGW progeny and σ^e2^ the residual variance extracted from the anova of the RI and GW elementary plots; broad-sense heritabilities were calculated as H^2^ = (σ^t2^-σ^e2^)/σ^t2^. V_WR = visual estimation of the presence of white rot in situ; V_TN = visual estimation of the presence of the total necrosis (degraded wood and white rot tissue) in situ; I_WR = proportion of white rot in % of trunk section area, measured by image analysis; I_TN = proportion of total necrosis in % of trunk section area, measured by image analysis; I_TA = area of the trunk section in cm^2^, reflecting trunk vigor and measured by image analysis.

| | Min. value | Median | Max. value | $\sigma_t^2$ | $\sigma_e^2$ | $H^2$ |
| --- | --- | --- | --- | --- | --- | --- |
| V_WR | 0.00 | 1.00 | 5.67 | 1.56 | 0.98 | 0.372 |
| V_TN | 1.00 | 3.67 | 8.33 | 2.42 | 1.83 | 0.241 |
| I_TA (cm<sup>2</sup>) | 201.30 | 324.80 | 505.90 | 2859,97 | 1249,09 | 0.563 |
| I_WR (%) | 0.00 | 0.55 | 16.15 | 0.0005 | 0.0003 | 0.412 |
| I_TN (%) | 4.70 | 14.04 | 41.58 | 0.0040 | 0.0019 | 0.507 |

To assess the relationship between measured variables, Pearson correlation coefficients were calculated. All variables are positively correlated (Table 2). As expected, relationship between visual scorings and image analyses for the same necrosis type are the strongest (r = 0.78 between V_TN and I_TN and r = 0.83 between V_WR and I_WR). Interestingly, a correlation was also observed between white rot and total necrosis, with correlation coefficients ranging from 0.48 to 0.57. Nevertheless, some genotypes, although displaying necrosis, did not show white rot (Figure S1). It is also noteworthy that a weak but significant positive correlation was detected between the trunk development and the proportion affected by necrosis, with r coefficients ranging from 0.34 to 0.43.

**Table 2.** Pearson correlation coefficients between all pairs of variables measured on RIxGW population. All the correlations are significant at p=0.001.

| V_TN | I_TA | I_WR | I_TN |  |
| --- | --- | --- | --- | --- |
| 0.56519 | 0.43203 | 0.82900 | 0.51836 | V_WR |
|  | 0.43269 | 0.50117 | 0.77682 | V_TN |
|  |  | 0.33744 | 0.35945 | I_TA |
|  |  |  | 0.48417 | I_WR |

### Gewurztraminer susceptibility to esca is governed by a single dominant factor located on grapevine chromosome 1

To decipher the genetic basis of the observed variations in white rot and total necrosis, a quantitative trait locus (QTL) analysis was performed in RIxGW population. To this end, we used genotyping-by-sequencing (GBS) data acquired on a set of 252 individuals of the progeny to establish two parental genetic maps. Both maps cover 19 linkage groups, corresponding to the 19 chromosomes of *V. vinifera*, and a high marker density, with an average distance of 0.1 cM between markers (Table 3, Figure S2). The female (RI) map includes 9 449 SNPs, with a total genetic length of 1239 cM. The male (GW) map has 9 427 SNPs covering 1175 cM.

**Table 3.** Main features of RI and GW parental maps.

| Linkage group | Riesling linkage map |  |  |  | Gewurztraminer linkage map |  |  |  |
| --- | --- | --- | --- | --- | --- | --- | --- | --- |
|  | Number of markers | Length (cM) | Average spacing (cM) | Max spacing (cM) | Number of markers | Length (cM) | Average spacing (cM) | Max spacing (cM) |
| 1 | 595 | 76.6 | 0.1 | 3.6 | 690 | 74.2 | 0.1 | 3.6 |
| 2 | 413 | 56.8 | 0.1 | 6.8 | 261 | 33.0 | 0.1 | 5.6 |
| 3 | 274 | 71.9 | 0.3 | 7.2 | 430 | 54.4 | 0.1 | 3.2 |
| 4 | 608 | 67.5 | 0.1 | 5.6 | 653 | 72.3 | 0.1 | 5.6 |
| 5 | 592 | 72.9 | 0.1 | 12.1 | 363 | 86.2 | 0.2 | 8.0 |
| 6 | 606 | 58.7 | 0.1 | 2.4 | 569 | 63.5 | 0.1 | 2.0 |
| 7 | 678 | 90.1 | 0.1 | 5.6 | 725 | 91.3 | 0.1 | 6.4 |
| 8 | 635 | 69.1 | 0.1 | 4.0 | 618 | 63.1 | 0.1 | 3.2 |
| 9 | 409 | 64.3 | 0.2 | 4.4 | 383 | 54.0 | 0.1 | 7.2 |
| 10 | 395 | 35.3 | 0.1 | 3.6 | 537 | 55.2 | 0.1 | 2.8 |
| 11 | 552 | 66.3 | 0.1 | 3.6 | 335 | 58.4 | 0.2 | 7.6 |
| 12 | 406 | 50.4 | 0.1 | 3.6 | 537 | 53.2 | 0.1 | 2.8 |
| 13 | 454 | 72.5 | 0.2 | 12.6 | 366 | 60.4 | 0.2 | 6.8 |
| 14 | 843 | 79.4 | 0.1 | 3.6 | 653 | 63.1 | 0.1 | 3.2 |
| 15 | 355 | 57.2 | 0.2 | 2.8 | 374 | 51.2 | 0.1 | 2.8 |
| 16 | 361 | 52.8 | 0.1 | 4.4 | 498 | 54.0 | 0.1 | 4.8 |
| 17 | 365 | 62.8 | 0.2 | 7.6 | 329 | 50.3 | 0.2 | 12.6 |
| 18 | 629 | 91.0 | 0.1 | 6.4 | 726 | 90.1 | 0.1 | 4.4 |
| 19 | 279 | 44.1 | 0.2 | 2.8 | 380 | 47.2 | 0.1 | 2.0 |
| **Overall** | **9449** | **1239.7** | **0.1** | **12.6** | **9427** | **1175.1** | **0.1** | **12.6** |

QTL detection was performed for all the measured traits. A single region located on chromosome 1 was detected on the GW parental map for all of the four traits describing necroses associated to esca (V_TN, I_TN, V_WR and I_WR) (Table S2; Figure 3). But no necrosis-related factors were identified on the RI parental map. This strongly suggests that white rot and total necrosis are both governed by a unique dominant factor which would be heterozygous in GW and which we have named *ENS1* for ‘Esca Necrosis Susceptibility 1’. The part of genetic variance explained by the variation of *ENS1* ranged from 14.6 % for I_WR to 51.1 % for V_TN. Noticeably, for each type of necrosis, the visual scoring appeared more efficient than image analysis for detecting QTL, both through the LOD score and through the part of variance explained by *ENS1*.

**Figure 3.**
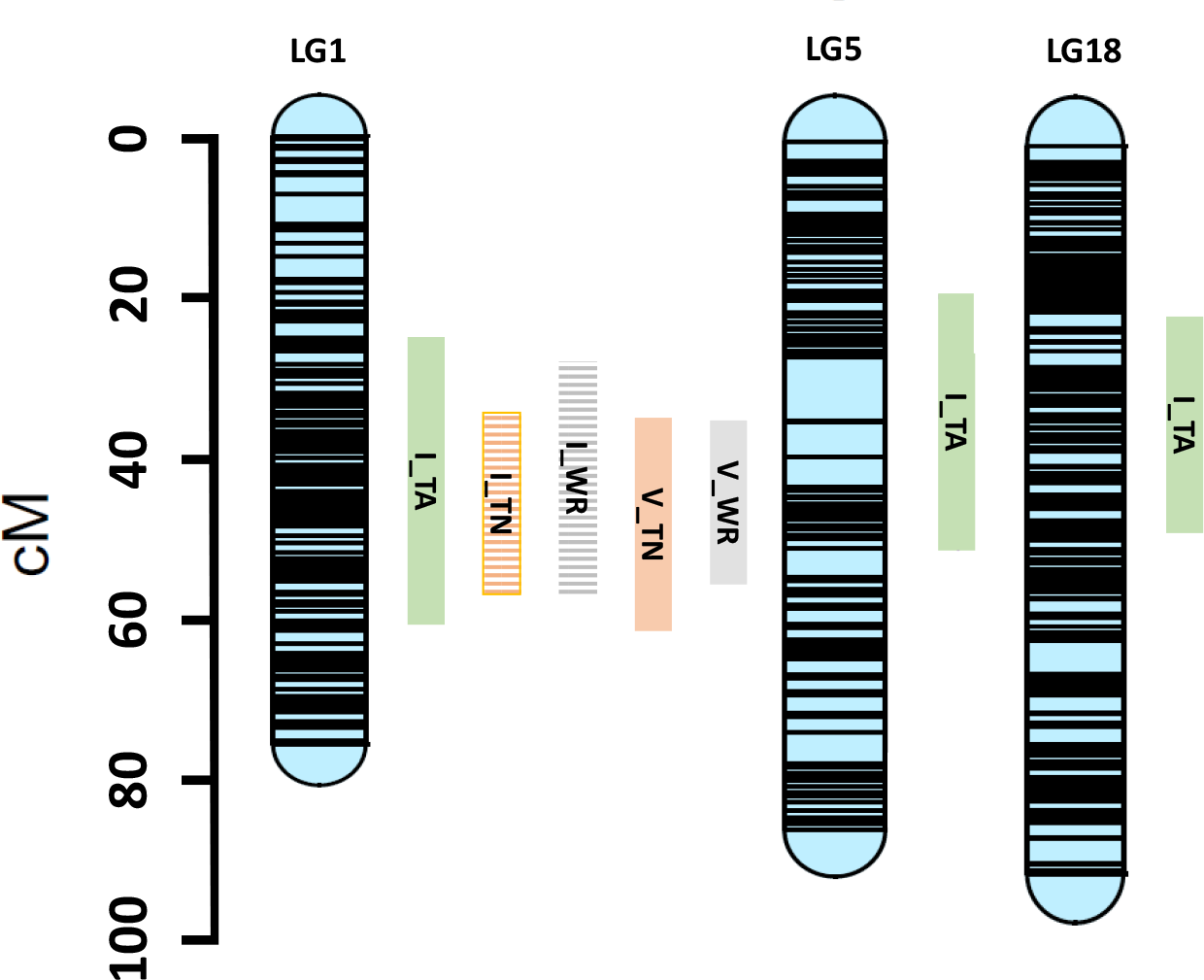
Location of the confidence intervals of QTL related to necrosis and trunk vigor detected on GW linkage map. V_WR = visual estimation of the presence of white rot in situ; V_TN = visual estimation of the presence of the total necrosis (degraded wood and white rot tissue) in situ; I_WR = proportion of white rot in % of trunk section area, measured by image analysis; I_TN = proportion of total necrosis in % of trunk section area, measured by image analysis; I_TA = area of the trunk section in cm^2^, reflecting trunk vigor and measured by image analysis.

Three QTL were detected for trunk development (I_TA), one on chromosome 1 on GW map, overlapping a region including *ENS1*, one on chromosome 5 on GW map and another one on chromosome 18 on both parental maps (Table S2; Figure 3).

In order to confirm the identification of *ENS1* discovered in Gewurztraminer, we used an alternative population derived from GW self-pollination (S1GW; 86 progeny) planted in an experimental design similar to the one used for RIxGW progeny, which was 20-year old (experiment B: e.p. of 3 plants; 1 e.p. for each S1GW descendant; 11 e.p. of GW7). Based on the results obtained with the RIxGW population, we estimated the susceptibility to esca necrosis in the S1GW population by visual scoring of necrosis. Genotyping was performed using a locus-specific Kompetitive Allele Specific PCR (KASP) marker designed in the *ENS1* region and named Chr1_10021151.

We first validated the Chr1_10021151 marker by comparing the effect of its allelic variation on a new subset of the RIxGW population (54 progeny) to the effect of a SNP at the same locus on the reference RIxGW population used to identify *ENS1*. Chr1_10021151 allowed to characterize genotype GW as heterozygote (XY) and RI as homozygous (YY). Despite the difference of size between both sets of RIxGW progeny, the effects revealed by the KASP marker were very similar to those revealed by the SNP at the same locus, for all the necrosis traits (Table 4), which confirmed that the KASP marker is appropriate to analyze the presence of *ENS1* on an alternative population.

**Table 4.** Validation of the Chr1_10021151 KASP marker. To validate the Chr1_10021151 KASP marker, the effect of its variation on the variables associated to necrosis and vigor measured on a RIxGW new subset was compared to the effect of a SNP of the *ENS1* region on the reference RIxGW population. For the SNP marker, A and H represent respectively homozygous recessive and heterozygous genotypes of the progeny. For the Chr1_10021151 KASP marker, YY and XY represent respectively homozygous recessive and heterozygous genotypes of the progeny.

|  | RlxGW reference population |  |  | New RlxGW subset |  |  |
| --- | --- | --- | --- | --- | --- | --- |
|  | H (mean) | A (mean) | P-value<br>(Kruskal-Wallis) | XY (mean) | YY (mean) | P-value<br>(Kruskal-Wallis) |
| V_WR | 1.68 | 0.93 | 2.164e-06 | 2.02 | 0.61 | 0.001649 |
| V_TN | 4.16 | 3.20 | 2.071e-07 | 5.13 | 3.53 | 0.001058 |
| I_TA (cm<sup>2</sup>) | 335.18 | 313.11 | 0.0007603 | 353.18 | 316.35 | 0.04334 |
| I_WR (%) | 1.87 | 0.89 | 0.0004223 | 3.25 | 1.03 | 0.009614 |
| I_TN (%) | 16.05 | 12.53 | 5.636e-06 | 21.25 | 13.55 | 0.0002158 |

The S1GW progeny segregated for both necrosis traits, in a range similar to that of RIxGW (Figure 4). Total necrosis visual scores ranged from 1 to 9 and white rot from 0 to 9. The Chr1_10021151 KASP marker allowed classifying the population individuals into 3 genotypes (XX:XY:YY), with no significant difference detected between the observed and expected Mendelian ratios (Table 5). Differences between genotypes are significant for both variables, V_TN and V_WR. Mean score comparison of the three genotypes allowed to confirm that the *ENS1* allele associated to susceptibility to esca necroses, and corresponding to the KASP marker X allele, is dominant (Table 5).

**Table 5.** Effect of the segregation of the Chr1_10021151 KASP marker on necrosis (V_TN and V_WR) in S1GW progeny. -: not applicable.

|  | Genotypes |  |  | Statistical tests |  |
| --- | --- | --- | --- | --- | --- |
|  | XX | XY | YY | P-value<br>(Kruskal-Wallis) | P-value<br>(Chi-square) |
| Number of genotypes | 16 | 50 | 20 | - | 0.2656 |
| V_WR mean | 4.16 | 4.19 | 2.43 | 0.0107 | - |
| V_TN mean | 5.24 | 4.54 | 3.75 | 0.0218 | - |

**Figure 4.**
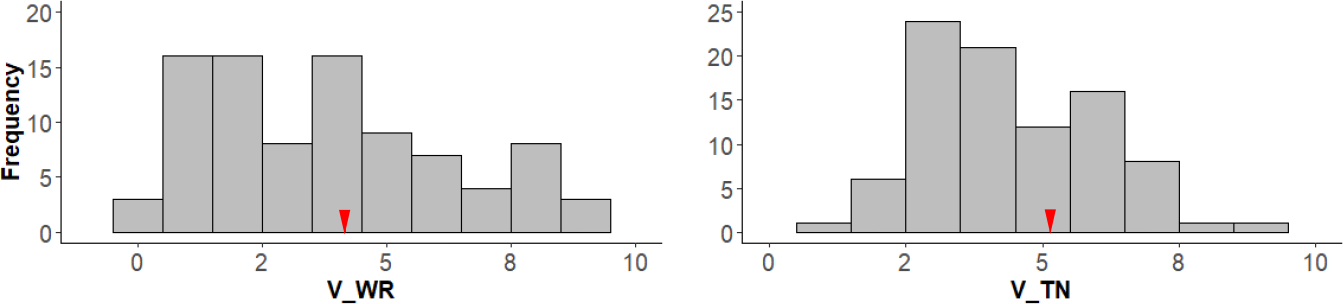
Distribution of necrosis parameters in the S1GW progeny. Mean value for the GW control is represented with a red arrow.

### Gewurztraminer susceptibility to esca is partly linked to trunk vigor

As mentioned previously, a partial link between, on the one hand, trunk section area, and on the other hand, total necrosis and white rot was observed in RIxGW progeny both at phenotypic level, with Pearson correlation coefficients ranging from 0.34 to 0.43 (Table 2) and at genetic level, with co-location of one QTL determining trunk section area with *ENS1* (Table S2; Figure 3).

To characterize this relationship, we performed a covariance analysis that decomposed the variances of variables related to necrosis (V_TN and V_WR) into three components: variance explained by the covariate I_TA, variance explained by ENS1, and the interaction between I_TA and ENS1 (Table 6). As expected, both allelic form at *ENS1* and I_TA had a significant effect on total necrosis and white rot. Interestingly, we observed an interaction between these two variables. The slope and the correlation coefficient associated to linear regression model between necrosis variables and trunk section area differed based on the *ENS1* genotype under consideration (Figure 5). For both V_TN and V_WR, the correlation was highly significant among individuals carrying the *ENS1* allele associated to susceptibility whereas the relationship was much weaker among individuals lacking the susceptibility-associated *ENS1* allele (Table S3).

**Table 6.**
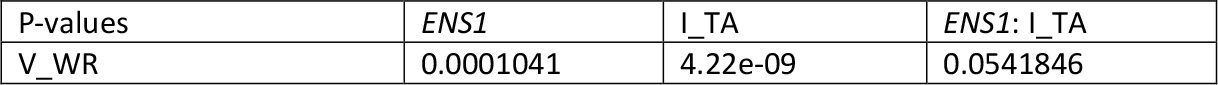
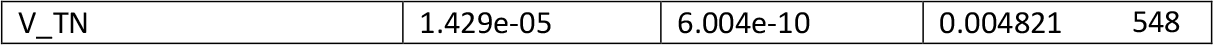
Covariance analysis of the effects of *ENS1*, trunk section area (I_TA) and their interactions on necrosis (V_TN and V_WR) in the reference RIxGW population.

| P-values | *ENS1* | I_TA | *ENS1*: I_TA |
| --- | --- | --- | --- |
| V_WR | 0.0001041 | 4.22e-09 | 0.0541846 |

|  |  |  |  |  |
| --- | --- | --- | --- | --- |
| V_TN | 1.429e-05 | 6.004e-10 | 0.004821 | 548 |

**Figure 5.**
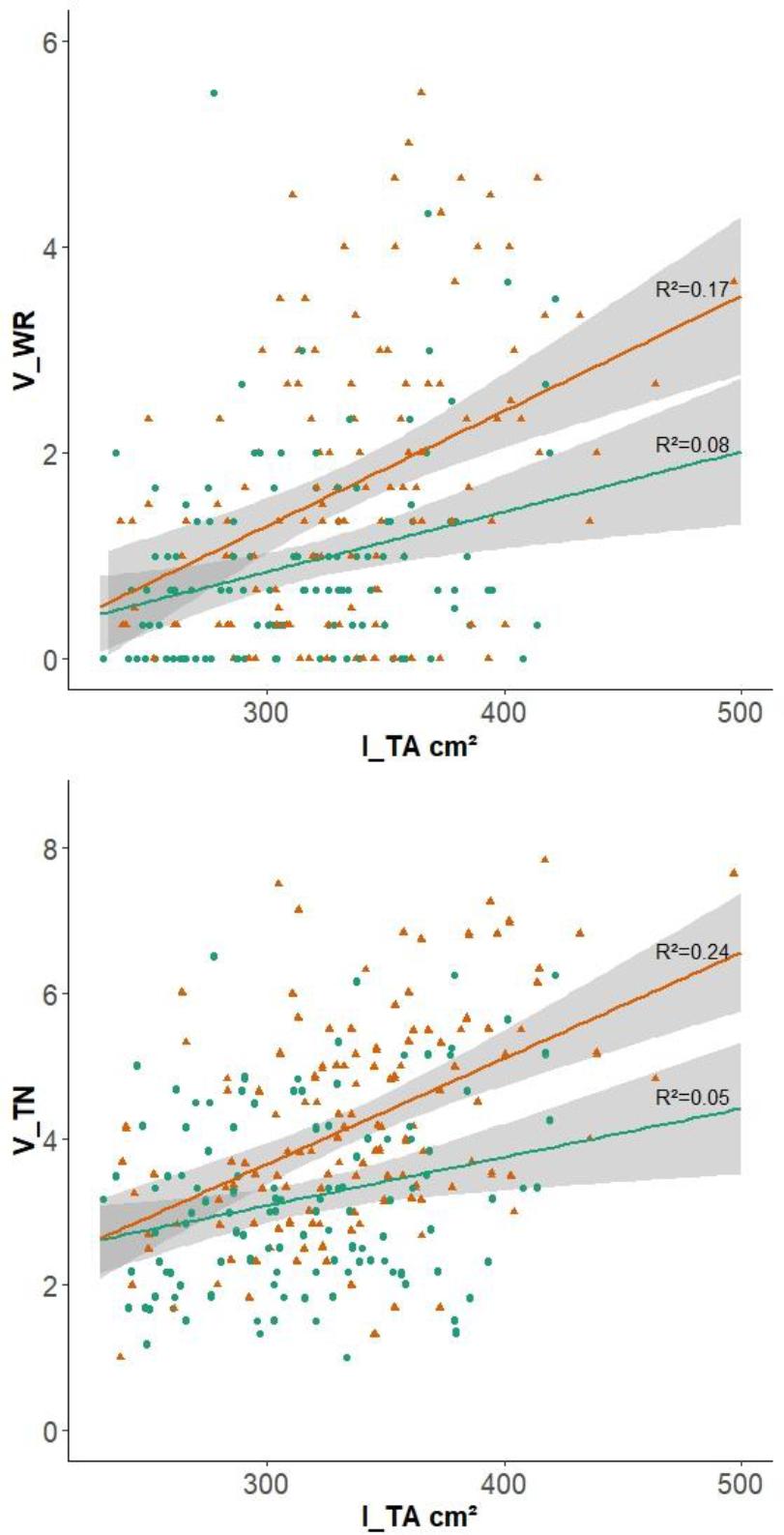
Scatter plots of total necrosis and white rot against vigor expressed by trunk section area according to the presence of *ENS1*. The data were recorded on the reference RIxGW population The lines represent linear regressions and the shaded areas their confidence intervals. In green, the individuals carrying at the chr11089995 SNP, located in the vicinity of the *ENS1*, the genotype (A) linked low susceptibility, and in red, those carrying the genotype (H) linked to high susceptibility.

## Discussion

Our study allowed us to identify, to our best knowledge, the first and, to date, the only instance of genetic factor involved in the limitation of necrosis associated to grapevine esca. Indeed, while differences in the frequency and incidence of esca have already been reported between grapevine varieties, no evidence of the genetic origin of these observations has been provided so far. Despite the importance of GTDs in general and esca in particular in vineyard decline, several reasons can be put forward to explain why it is so difficult to identify genetic factors that could provide effective and sustainable solutions to these diseases, particularly through breeding of new grapevine tolerant varieties. A first reason is related to the complexity of the etiology and the multifactorial nature of the symptomatic expression of esca, making it difficult to set up a bioassay under controlled conditions capable of reproducing the differences observed in the vineyard. A second obstacle is that, in addition to the etiology complexity, the symptoms in the vineyard can take a long time to appear. A third reason is that some of the effects of the disease, such as necrosis, are internal and therefore not easily detectable. For all these reasons, we have chosen to conduct this study on plants grown in the vineyard and at least 16 years old. The choice of the plant material was also crucial and aimed at optimizing the chance of observing segregation, based on the prior observation that Riesling and Gewurztraminer showed a differential response to esca in the vineyard^9,35^. The evaluation of a phenotype unambiguously linked to a severe form of esca was the trickiest point to achieve, for which we implemented a destructive method by longitudinally sectioning the trunk of the plants in order to directly observe the internal symptoms.

A wide variation in the proportion of the grapevine trunk affected by necrosis has been observed and associated to a single locus located on grapevine chromosome 1, which we have named *ENS1*. The favorable allele of *ENS1*, limiting the development of necrotic trunk tissue, is deduced to be recessive. This result is of strategic importance given that esca is a major concern for the wine-growing sector worldwide, leading to vineyard degeneration, and that no sustainable and environmentally friendly method of control exists so far, despite the many efforts by various research groups over decades^6,14^. More than 600 genes were counted in the QTL confidence interval which makes it difficult to identify a short list of candidate genes. However it is interesting to note the co-location of *ENS1* with VvWRKY2, a transcription factor described as possibly playing a role in tolerance to necrotrophic fungal pathogen, lignin biosynthesis and xylem development^36,37^.

Nevertheless, the two critical key points of the approach we have used remains the long experimental duration and the destructive phenotyping. Given the observed complexity of the interactions between biotic and abiotic factors in the expression of esca, producing symptoms in a laboratory model is currently challenging. Such development is particularly complicated by the number of factors (fungal species potentially involved and experimental conditions adapted to disease expression), or even combinations of factors, to be tested. In this context, the characterization of the plant material resulting from our study could be used to establish the basis for a rapid bioassay. The possibility of discriminating genotypes according to their genetic predisposition to develop necrosis in the vineyard from the segregation of susceptibility alleles provides a reference sample better adapted to the development of a phenotyping methods in controlled inoculation conditions than a collection of varieties that do not always show consistent field performance. It would then be possible to better specify the role of the different putative pathogens with respect to the observations made in the field.

Necrosis is directly related to the development of esca and vine mortality. It is therefore crucial to be able to assess necrosis in a grapevine trunk in further genetic studies. Unfortunately, internal necroses are difficult to observe without cutting the plant trunk. Indeed, although the characterization method used in our study was effective in quantifying internal trunk necrosis, it required a feasible destructive sampling. If one favors the use of plant material that is old enough to allow the development of disease symptoms in the vineyard, as in the case of our study, it would be critical to develop instruments capable of assessing internal esca damage in the vineyard in a non-destructive procedure. Non-destructive measurement systems using magnetic resonance imaging (MRI) and X-ray tomography are under development^22,38,39^ and will certainly be able to help with the implementation of non-destructive studies on old vines in the future. Such non-destructive measurement systems will also be important to study the dynamics of the development of internal necroses over the time.

This study also suggests that there is a link between trunk vigor and necrosis due to esca both at phenotypic level, through a positive correlation, and at genetic level, through co-location of QTLs. The decomposition of the correlation clearly showed that a part of necrosis is determined by the interaction between *ENS1* and trunk vigor. A difference in terms of norm of reaction to disease infection depending on the presence of *ENS1* is one of the expressions of the observed link between trunk vigor and necrosis due to esca (Figure 5). The most tolerant individuals react very weakly to the variation of vigor whereas susceptible individuals react significantly. The observation of this difference in norms of reaction seems consistent with the link between vine vigor and esca development that has often been described in previous studies^18–20^, the most susceptible individuals displaying this link, while the most resistant ones do not. Furthermore, trunk vigor was determined in this study by three QTLs, one on chromosome 1 linked to *ENS1* and the other on chromosomes 5 and 18 not linked to regions involved in the variation of esca necrosis. This suggests that such a potential link between vigor and necrosis is only partial, as not all the variation in necrosis can be explained by a variation in vigor. Regarding the overlap of the QTL determining trunk vigor and esca necrosis on chromosome 1, three hypotheses can be proposed: a genetic co-location of two functionally independent factors, specific to necrosis on the one hand and to vigor on the other; a gene with a pleiotropic effect on both, vigor and esca necrosis, traits; a physiological relationship between vigor and necrosis. While the first two situations are perfectly plausible, the last one seems more difficult to explain because the part of the variation in vigor determined by the QTLs on chromosomes 5 and 18 was not linked to necrosis.

The relationship between total necrosis and white rot was strong, both at the genetic and phenotypic levels. This is in line with a sequential development of woody tissue degradation already described, moving from healthy wood to dead wood, then from dead wood to white rot^28,31^. These observations form the basis of recommended viticultural practices to limit the development of wood diseases, such as respectful pruning to limit the formation of dead wood and trunk surgery to eliminate white rot^30,31^.

To conclude, even if the identification of *ENS1* alone will not solve the esca issue, this discovery is nonetheless a first step towards a genetic solution. Indeed, our study proves that it is possible to associate a genetic factor with variations in susceptibility to esca observed in the vineyard. In so doing, this finding demonstrates the potential existence of a source of resistance or tolerance to esca in the diversity of *Vitis* that remains to be identified and, thus, paves the way and encourages future genetic studies of greater scope.

## Supplementary data

Supplementary information is provided as a pdf document.

## Acknowledgments

We are grateful to C. Rustenholz and A. Velt for providing Gewurztraminer sequences used to develop the KASP marker, to the Vignes Vivantes association for performing the vine trunk sections and the Gentyane platform for KASP analyses. We would like to thank P. Mestre for his careful reading which helped improve the manuscript. We also thank our colleagues of the Unité Expérimentale Agronomique et Viticole INRAE-Colmar for maintaining the vineyard plots and E. Chedid for her advice on QTL analysis.

## Author contributions

GA and DM designed the research, interpreted the data and wrote the manuscript. GA carried out phenotyping and statistical analysis. EP managed the KASP marker development. VD set up and monitored the experimental plots. GB generated DNA libraries. KA established the genetic linkage maps and carried out statistical analysis. ED and GA carried out QTL detection. DM supervised the study. All authors have contributed to the revision of the manuscript.

## Data availability

The data underlying this article are available in the article, in the supplementary information files and in the online supplementary material at https://doi.org/10.57745/BWY9QK. 00

