## Supplemental Tables for "Discovery of a locus associated with susceptibility to esca dieback in grapevine"

**Table S1. Proportion of the trunk affected by necroses in control varieties, Riesling and Gewurztraminer, in experiment A**. Necrosis parameters (V_WR, V_TN, I_WR and I_TN) and trunk section area (I_TA) were assessed either by visual scoring (V_ variables) or imaging (I_ variables). Kruskal-Wallis test: **, ***: significant differences at P = 0.01 and P = 0.001, respectively; ns: not significant at P = 0.05.

| | GW | RI | P-value |
|---|---|---|---|
| V_WR | 1,47 | 1,06 | 0.323 ns |
| V_TN | 4,25 | 3,3 | 0.005553 ** |
| I_TA (cm²) | 396,77 | 332,05 | 8.589e-06 *** |
| I_WR (%) | 1,38% | 1,22% | 0.4428 ns |
| I_TN (%) | 16,29% | 14,66% | 0.09327 ns |

**Table S2. QTL analysis for the necrosis traits (V_WR, V_TN, I_WR and I_TN) and trunk section area (I_TA) from RI and GW parental maps.** CI low and CI high columns represent the limits of the QTL confidence interval, $R^2$, the part of total variance explained by the QTL and Vq/Vg, the part of genetic variance explained by the QTL.

| Map | Variable | LOD threshold at p=0.05 | Nearest marker | Chr | CI low | Peak pos. | CI high | LOD score | P-val | $R^2$ | $H^2$ | Vq/Vg |
|---|---|---|---|---|---|---|---|---|---|---|---|---|
| GW | V_TN | 3.15 | chr1_16184684 | 1 | 53.59 | 34.9 | 60.3 | 7.09 | 0.000 | 12,29% | 0,240 | 51,07% |
| GW | I_TN | 3.06 | chr1_10284333 | 1 | 44.06 | 34.1 | 56 | 5.92 | 0.000 | 10,38% | 0,507 | 20,46% |
| GW | V_WR | 2.99 | chr1_11089995 | 1 | 45.65 | 34.9 | 54.4 | 5.28 | 0.002 | 9,31% | 0,372 | 25,00% |
| GW | I_WR | 2.83 | chr1_7514826 | 1 | 38.11 | 27.8 | 56 | 3.35 | 0.015 | 6,01% | 0,412 | 14,58% |
| GW | I_TA | 3.03 | chr18_6620584 | 18 | 28.58 | 20.6 | 46.8 | 3.68 | 0.009 | 6,57% | 0,563 | 11,67% |
| GW | I_TA | 3.03 | chr5_6897497 | 5 | 43.74 | 19.4 | 50.9 | 3.5 | 0.017 | 6,26% | 0,563 | 11,12% |
| GW | I_TA | 3.03 | chr1_10564972 | 1 | 44.86 | 24.6 | 59.1 | 3.14 | 0.037 | 5,65% | 0,563 | 10,03% |
| RI | I_TA | 3.03 | chr18_12839502 | 18 | 51.24 | 7.1 | 58.8 | 3.92 | 0.007 | 6,99% | 0,563 | 12,41% |

**Table S3. Correlations between necrosis parameters and trunk section area measured according to the genotype at *ENS1* locus.** ). *, **, ***: significant differences at P = 0.05, P = 0.01 and P = 0.001, respectively.

| *ENS1* allele | V1 | V2 | Pearson's $R^2$ | P-value |
|---|---|---|---|---|
| A | I_TA | V_TN | 0.050 | 0.01409* |
| A | I_TA | V_WR | 0.084 | 0.00134** |
| H | I_TA | V_TN | 0.245 | 2.442e-09*** |
| H | I_TA | V_WR | 0.172 | 1.027e-06*** |

**Table S4. Primers designed for the Chr1_10021151 KASP marker.**

| Type | Name | Sequence |
|---|---|---|
| Allele-specific | chr1:10021151-10021450_1 | GAAGGTGACCAAGTTCATGCTTACGTTCATTTCTACATGTGCTACCT |
| Allele-specific | chr1:10021151-10021450_2 | GAAGGTCGGAGTCAACGGATTACGTTCATTTCTACATGTGCTACCA |
| Common | chr1:10021151-10021450 | AACCTTTATCGTGTAATAGTGATGGAAGAA |
