## Supplemental Figures1&2 for "Discovery of a locus associated with susceptibility to esca dieback in grapevine"

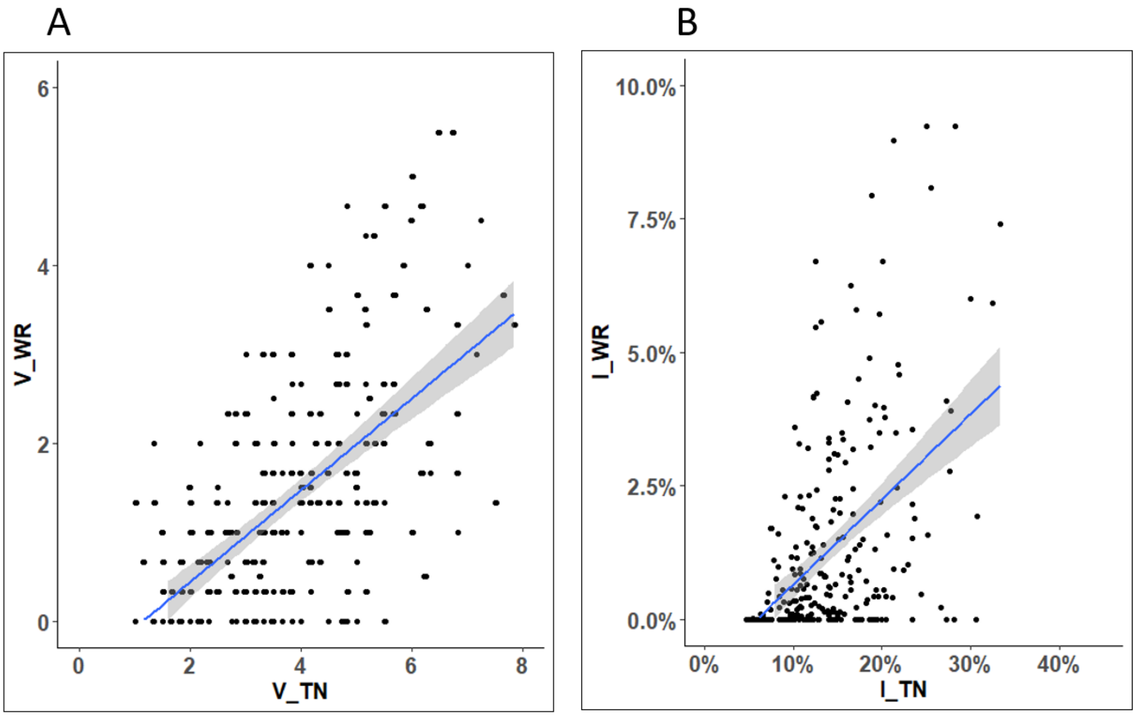


**Figure S1 : Scatter plots of white rot against total necrosis scored visually (panel A) and measured by imaging (panel B).** The blue lines represent linear regressions and the shaded areas their confidence intervals.

A

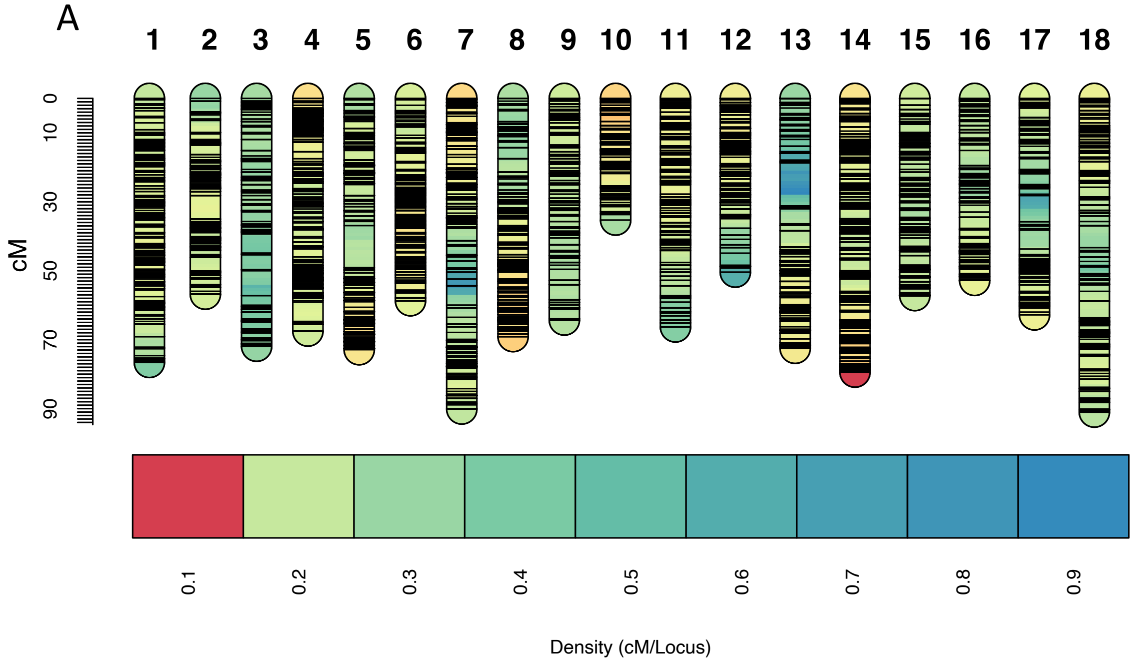


B

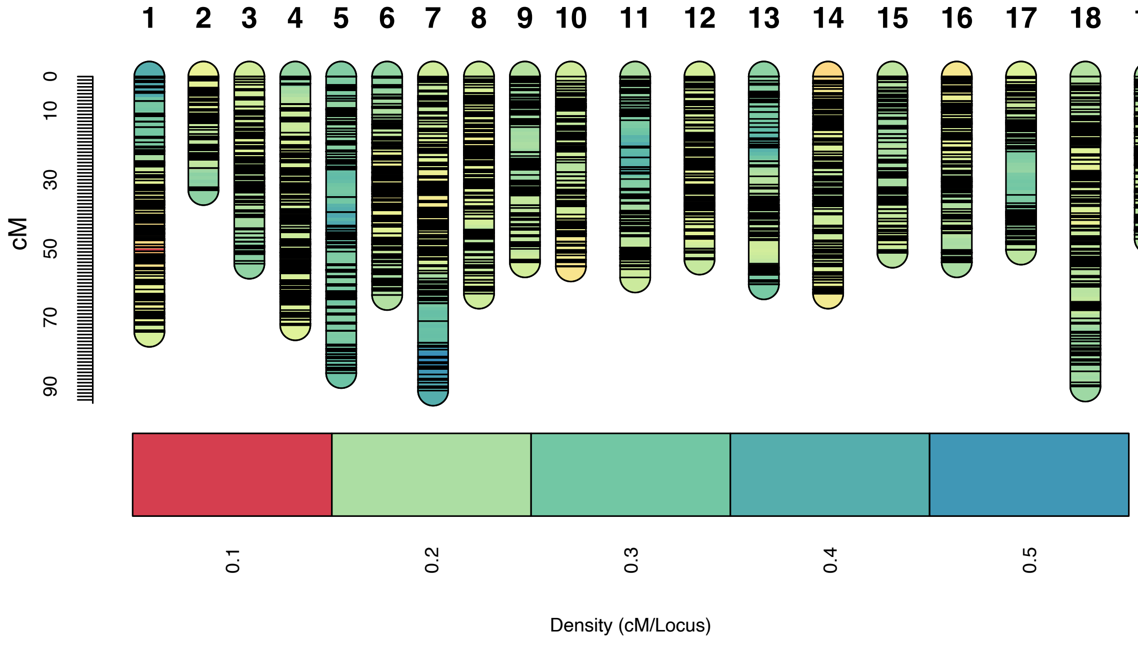


**Figure S2 : Riesling (A) and Gewurztraminer (B) parental maps.** Diagrams represent linkage groups. Black lines indicate marker position. The y–axis indicates genetic distance in centimorgans (cM). The color scale indicates marker density in cM/locus.
