## Supplemental Figure3 for "Discovery of a locus associated with susceptibility to esca dieback in grapevine"

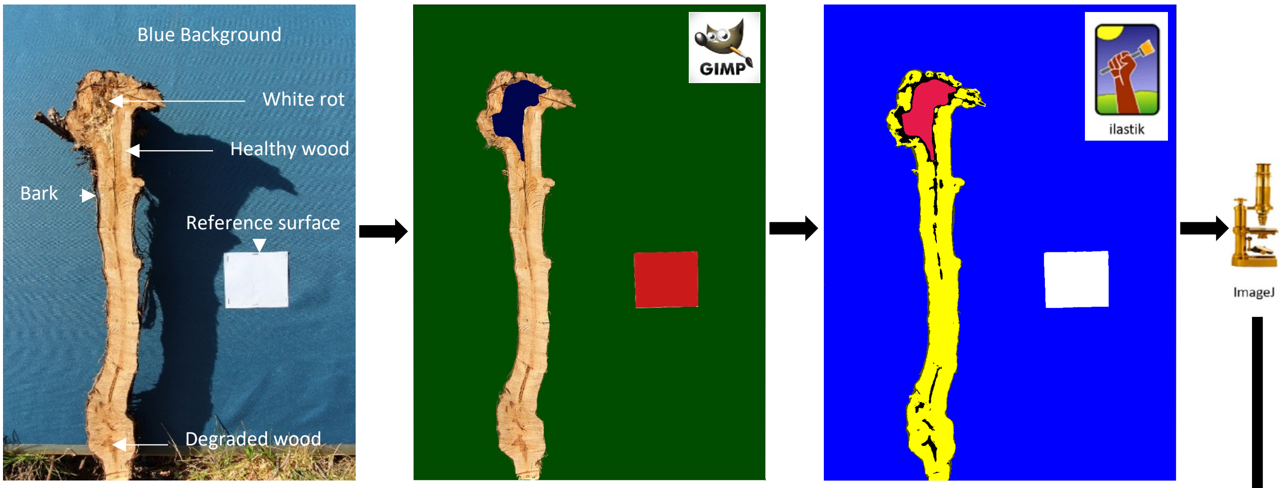


| Illastik colour | Blue Background | White Reference surface | Red White rot | Yellow Healthy wood | Black Degraded wood | Green Bark |
|---|---|---|---|---|---|---|
| **Channel (C)** | 1 | 2 | 3 | 4 | 5 | 6 |
| **Number of pixel** | 3 662 093 | 87 957 | 44 715 | 301 636 | 42 211 | 28 888 |

**Figure S2. Workflow for quantifying white rot and total necrosis extent and trunk section area (I_TA) by imaging and determination of necrosis parameters (I_TN, I_WR).** This example illustrates how the data used for the calculation of three variables (I_TA=(C3+C4+C5)x100/C2; I_WR=C3x100/(C3+C4+C5) ; I_TN =(C3+C5)x100/(C3+C4+C5)) are generated.
